# Proinsulin C-peptide is a major source of HLA-DQ8 restricted HIPs recognized by human Islet-Infiltrating CD4^+^ T cells

**DOI:** 10.1101/2024.05.09.593303

**Authors:** Pushpak Bhattacharjee, Miha Pakusch, Matthew Lacorcia, Eleonora Tresoldi, Alan F. Rubin, Abby Foster, Laura S. King, Chris Chiu, Thomas W.H. Kay, John A. Karas, Fergus J. Cameron, Stuart I. Mannering

## Abstract

Type 1 diabetes (T1D) is an autoimmune disease that develops when T cells destroy the pancreatic insulin-producing beta cells that reside in the pancreatic islets. Immune cells, including T cells infiltrate the islets and gradually destroy the beta cells. Human islet-infiltrating CD4^+^ T cells recognize peptide epitopes derived from proinsulin, particularly C-peptide. Hybrid Insulin peptides (HIPs) are neoepitopes formed by the fusion of two peptides derived from beta-cell granule proteins and are known to be the targets of pathogenic CD4^+^ T cells in the NOD mouse and human islet-infiltrating CD4^+^ T cells. Proinsulin is widely recognized as a central antigen in T1D, but its role in forming HIPs is unclear. We developed a method to functionally screen TCRs derived from human islet-infiltrating CD4^+^ T cells and applied this to the identification of new proinsulin-derived HIPs. We generated a library of 4,488 candidate HIPs formed by fusion of proinsulin fragments and predicted to bind to HLA-DQ8. This library was screened against 109 islet-infiltrating CD4^+^ T-cell TCRs isolated from four organ donors who had T1D. We identified 13 unique HIPs recognized by 9 different TCRs from two organ donors. HIP specific T-cell avatars responded specifically to a peptide extract from human islets. These new HIPs predominantly stimulated CD4^+^ T-cell proliferation in PBMCs from people with T1D in contrast to HLA-matched controls. This is the first unbiased functional, islet-infiltrating T-cell based, screen to identify proinsulin derived HIPs. It has revealed many new HIPs and a central role of proinsulin C-peptide in their formation.

**SUMMARY:** Type 1 diabetes is an autoimmune disease caused by T cells destroying the pancreatic insulin-producing beta cells. The antigens/epitopes seen by disease promoting CD4^+^ T cells are poorly understood. Hybrid insulin peptides (HIPs) are a new class of CD4^+^ antigen recognized by pathogenic NOD mouse CD4^+^ T cells. In humans very few HIPs recognized by human islet-infiltrating CD4^+^ T cells are known. We show that proinsulin HIPs are recognized by human islet-infiltrating CD4^+^ T cells from T1D donors and describe 13 new HIPs formed by fusion of proinsulin peptides. This work shows that proinsulin, particularly C-peptide, is a major contributor to the pool HIPs recognized by human islet-infiltrating CD4^+^ T cells and are therefore central to autoimmunity in T1D.

## Introduction

Type 1 diabetes (T1D) is a chronic, incurable, autoimmune disease caused by the T-cell mediated destruction of the pancreatic, insulin-producing, beta-cells [1]. This leads to insulin deficiency and dysregulation of glucose metabolism [2]. Genetic risk of developing T1D is strongly associated with the HLA class II, specifically the haplotypes: HLA-DR3-DQ2 (HLA-DRB1*03:01-DQA*05:01-DQB*02:01) and HLA-DR4-DQ8 (HLA-DRB1*04:01; DQA*03:01, DQB*03:02) [3]. Of these HLA-DR4-DQ8 haplotypes give the highest risk of developing T1D [3, 4].

The pathogenicity of a CD4^+^ T cell cannot be directly demonstrated in human studies, so we developed methods to isolate and characterised CD4^+^ T cells from the pancreatic islets of T1D organ donors [5]. We [5], and others [6-9], have shown that these cells are strongly implicated in the immune pathogenesis of T1D based on their presence at the site of autoimmune beta-cell destruction, their antigen specificity and HLA restriction. Based on the genetic associations outlined above, CD4^+^ T cells that recognize beta-cell antigens presented by DR4-DQ8 or DR3-DQ2 are more strongly implicated in autoimmune beta-cell destruction [5, 6, 8]. Insulin and its precursor proinsulin have emerged as important targets of autoimmune responses in T1D [10, 11]. Specifically, C-peptide, which is excised from proinsulin, has emerged as major antigenic target of human islet-infiltrating [5, 8] and peripheral blood CD4^+^ T cells [12].

CD4^+^ T-cell responses against neoepitopes formed by the modification of native proteins explain, how immune tolerance to otherwise healthy cells and tissues is lost [13, 14]. CD4^+^ T-cell responses to several neoepitopes have been associated with human T1D [14-17]. Hybrid insulin peptides (HIPs) [18] are neoepitopes that form by the fusion of two peptide fragments. CD4^+^ T-cells specific for HIPs are pathogenic in the NOD mouse model of human T1D [18]. BDC2.5 T cells that are specific for HIPs formed by the fusion of C-peptide and chromogranin-A transfer diabetes to NOD mice [18, 19].

CD4^+^ T-cell responses to HIPs are increasingly implicated in the immune pathogenesis of human T1D. We [18], and others [7, 20] have shown that HIP specific CD4^+^ T-cells infiltrate the pancreatic islets of organ donors who had T1D. The presence of HIP specific, HLA-DQ8 restricted CD4^+^ T cells within the pancreatic islets of people who had T1D strongly suggests that they play a causative role in the development of T1D. Studies using peripheral blood mononuclear cells (PBMCs) from people with and without T1D revealed that many of the original 12 HIPs identified by Delong et al [18] were capable of stimulating CD4^+^ T-cell responses in PBMC from people with T1D [21, 22]. Arribas-Layton et al [23] used HLA-DR4 tetramers and peptide binding assays to identify six HLA-DR4 (DRB1*04:01) restricted HIPs recognized by peripheral blood derived CD4^+^ T cells from people with T1D. Responses to HIPs arise early in the immune pathogenesis of T1D. CD4^+^ T-cell responses to HIPs have been reported to be detectable, in the peripheral blood, prior to the onset of clinical T1D [24]. Autoantibodies specific extended HIP sequences formed by the fusion of fragments of proinsulin with IAPP2 peptides were detected in the serum of individuals before insulin autoantibodies could be detected [25].

Understanding the immune biology of HIPs in T1D faces two related challenges [26]. First, since HIPs are formed by the fusion of two peptide fragments there is a very large number of candidate HIPs that can potentially form [26] making it very difficult to identify new HIPs. Second, it is difficult to link a CD4^+^ T-cell responses against a HIP to the pathogenesis of human T1D. To address these challenges we devised a high-throughput functional screening assay to identify proinsulin derived, HLA-DQ8 binding HIPs and. To accommodate the large number of potential HIPs we established an *E. coli* library of candidate, proinsulin derived, HLA-DQ8 binding HIPs. To ensure that we identified HIPs relevant to the immune pathogenesis of human T1D we used islet-infiltrating CD4^+^ T-cells to screen the HIP library and identify HIPs recognized by these cells. We reasoned that identifying HIPs formed from proinsulin, presented by HLA-DQ8 and recognized by human islet-infiltrating CD4^+^ T cells would of particular relevance to the pathogenesis of human T1D. Using this approach, we identified 13 new HIPs recognized by human islet infiltrating CD4^+^ T cells. Analysis of the sequence of these HIPs revealed that they all incorporated fragments of proinsulin C-peptide, indicating that C-peptide is an important contributor to T1D associated human HIPs.

## Results

### Generation of a panel of human islet-infiltrating CD4 T-cell avatars

Previously we have shown that two human islet-infiltrating CD4^+^ T-cell clones recognized a HIP formed by the fusion of C-peptide with IAPP2 [18]. Given the importance of HIPs in human CD4^+^ T-cell responses associated with T1D, we set out to systematically identify new HIPs. Specifically, we wanted to identify novel HIPs that were recognized by human islet-infiltrating CD4^+^ T cells. To achieve this, we generated and screened 109 T-cell avatars (Supplementary Figure 1). Each avatar expressed the TRAV and TRBV derived from a single CD4^+^ T cell isolated from one of our four organ donors who suffered from T1D. Two donors had T1D for 3 or 4 years, while the other two had T1D for 13 and 19 years (Supplementary Table 1).

### Bacterial-based antigen screening identifies new HIPs

We narrowed our search to HIPs that could form by the fusion of peptides derived from proinsulin (86 amino acids). We generated, *in silico*, a library of the 9,409 HIPs that could theoretically be formed by fusion of any two 12mer fragments of human proinsulin (Supplementary Figure 2A). We then filtered these candidate HIPs to select unique sequences which were within the top 21% of predicted HLA-DQ8 (HLA-DQA1*03:01; DQB1*03:02) binding affinities. This led to a list of 4,488 12mer candidate, proinsulin derived HIPs that predicted to bind strongly to HLA-DQ8. Hence, we screened 489,192 (109 x 4,488) combinations of T-cell avatars and candidate HIPs. Oligonucleotides encoding these candidate HIPs were synthesized cloned into the expression vector pKE-1 [27] and used to transform *E. coli*. The library of 4,488 candidate HIPs was divided into 136 pools each comprising 34 candidate HIP sequences. To validate the library, a sample was sequenced and the abundance of each target sequence was determined (Supplementary Figure 2B). These data show that >97.5% of the target sequences were present in the library at an abundance of >26 TPM.

To identify novel proinsulin HIPs, we first optimized the workflow for detecting CD4^+^ T-cell responses by screening *E. coli* pools (Summarized in supplementary Table 2). The candidate HIP library was screened in three steps. For the first round of screening 34 ‘super-pools’ were prepared by combining groups of 4 of the 136 pools. This made it feasible to screen all 4,488 candidate HIPs could be against 109 T-cell avatars (Figure 1A). The results from screening one T-cell avatar (A_4_8) at each of the three steps is shown in Figure 1. In this case, super-pool 18 stimulated a strong response from avatar A_4_8. The second step was to screen the avatars that responded to a super-pool against the same super-pool and each of the four component pools (Figure 1B). Avatar A_4_8 responded to a HIP expressed in pool 71. Finally, the T-cell avatars that responded to each pool were tested against 40-80 individual *E. coli* clones isolated from the stimulating pool (Figure 1C). Colony 29 was the only colony screened that activated the TCR expressed in avatar A-4_8. The HIP sequences were determined by sequencing the plasmids isolated from the single stimulating colonies. After screening 109 avatars we had identified nine human islet-infiltrating T-cell avatars, six from Donor A and three from Donor K, that responded to 13 unique candidate HIPs. These donors had had T1D for 3 or 4 years. In contrast, no responses to any candidate HIPs were detected in the T-cell avatars isolated from the pancreatic islets of Donors I and J, who had had T1D for 13, or 19 years respectively (Supplementary Table 5). No correlation was seen between the abundance of the HIP sequences in the *E. coli* library and T-cell avatar responses (Supplementary Figure 2B) indicating that candidate HIPs could be identified even when they were relatively rare in the library.

**Figure 1.**
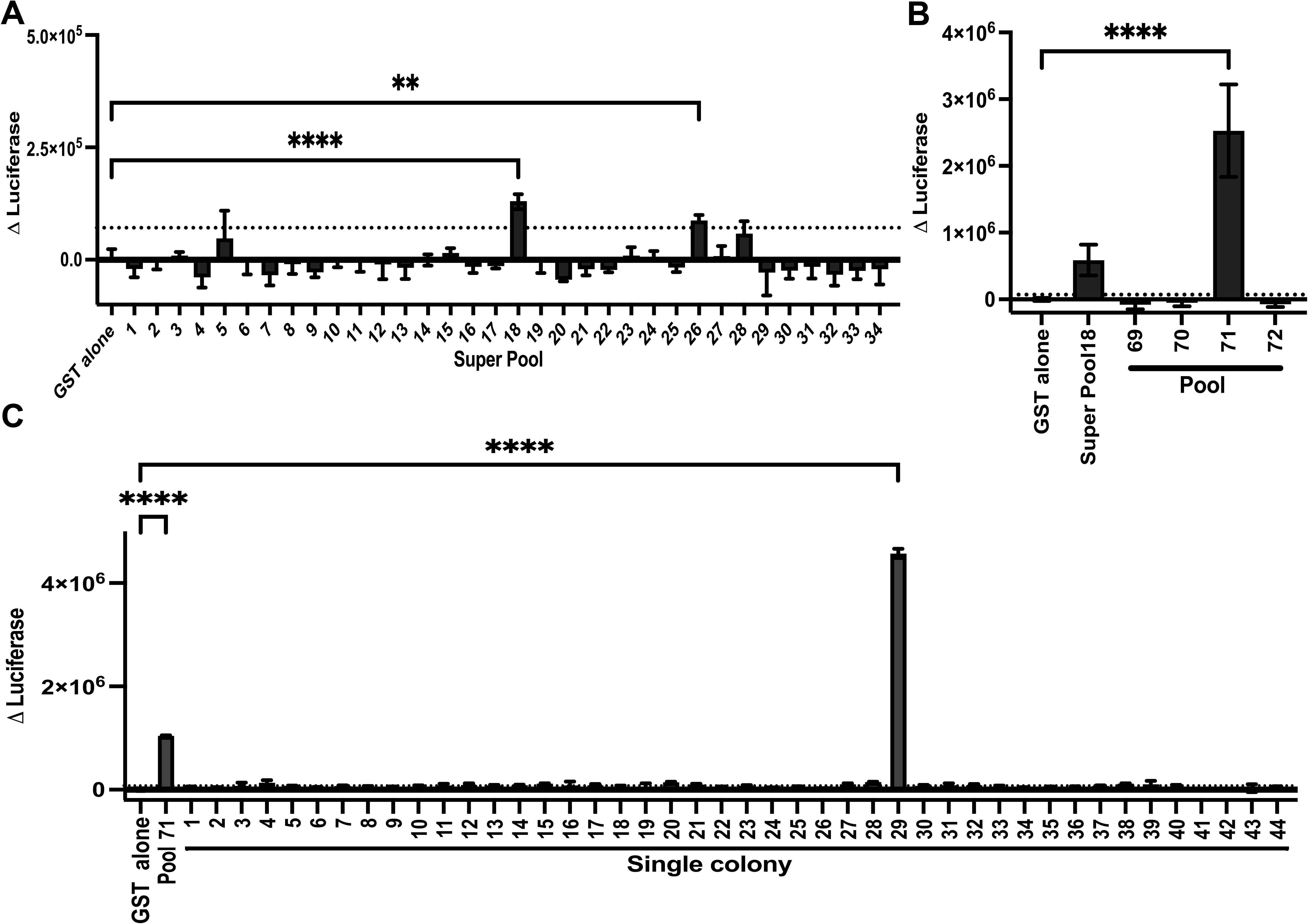
Functional screening of the candidate HIP library, expressed in E. coli, reveals new HIPs. An example of the screening of one T-cell avatar (A_4_8) is shown. (A) The HIP ‘super-pools’ were tested against the T-cell avatar A_4_8. (B) HIP ‘super-pools’ 18 and the four pools it comprises (pools: 69, 70, 71 and 72) were tested separately. (C) Forty-four *E. coli* colonies isolated from HIP Pool 71 were tested against the A_4_8 T-cell avatar. Antigen-specific responses were measured by luciferase activity in triplicate, except for the colony screening which was done in duplicate. The Δluciferase was calculated by subtracting the mean of the GST only plasmid-containing bacteria (GST alone) T-cell avatar, APC treatments from all other readings. The bars represent the mean of Δluciferase +/- SD. The dotted line is the GST alone treatment plus 2 x standard deviation (2SD) represents the threshold for a positive response. Statistical significance was determined using one-way ANOVA and corrected for multiple comparisons using Dunnett statistical hypothesis testing and defined as *p<0.05, **p<0.01, ***p<0.001, ****p<0.0001.

### Validating candidate HIP responses

To confirm that the T-cell avatars respond to the HIPs identified we synthesized peptides with the same sequence of each candidate proinsulin HIP (Supplementary Table 3). These peptides were used in titration experiments to measure their potency in stimulating the appropriate T-cell avatars. In all cases the responses to HIPs expressed in *E. coli* were confirmed with synthetic peptides (Figure 2). Some avatars recognized multiple HIPs, the avatar expressing the TCR A_4_2, responded to eight HIP sequences (Figure 2A), the most of any avatar. Two avatars (A_4_5, K_4_161 Figure 2B, 2H) responded to three HIPs and one (K_4_207, Figure 2I) responded to two HIPs. LogoPlots revealed that the C-terminal fragment remained the same, but TCR recognition could tolerate several sequences on the N-terminal side of the HIPs (Figure 2J-M). Conversely several HIPs stimulated more than one T-cell avatar. HIP2 and HIP4 stimulated T-cell avatars from Donor A (A_4_2) and Donor K (K_4_207). HIPs 1, 5, 6 and 7 were able to stimulate two (HIP1, HIP5), or three (HIP6, HIP7) Donor A T-cell avatars. Primary CD4^+^ T-cell clones from Donor A had previously shown to recognize epitopes derived from C-peptide [5, 18]. Three Donor A TCRs that we identified as being able to recognize proinsulin HIPs (A_4_2, A_4_7, A_4_8) also recognize epitopes derived from C-peptide [5] and two were previously shown to respond to a C-peptide-IAPP2 HIP [18] (Supplementary Table 6). However, using T-cell avatars, which are less sensitive, we detected very weak responses to full-length C-peptide at 20μM, the highest concentration tested from these avatars. In contrast, the HIPs were very potent agonists and responses could be detected with as little as <10-100 nM in the same assays (Supplementary Figure 3).

**Figure 2.**
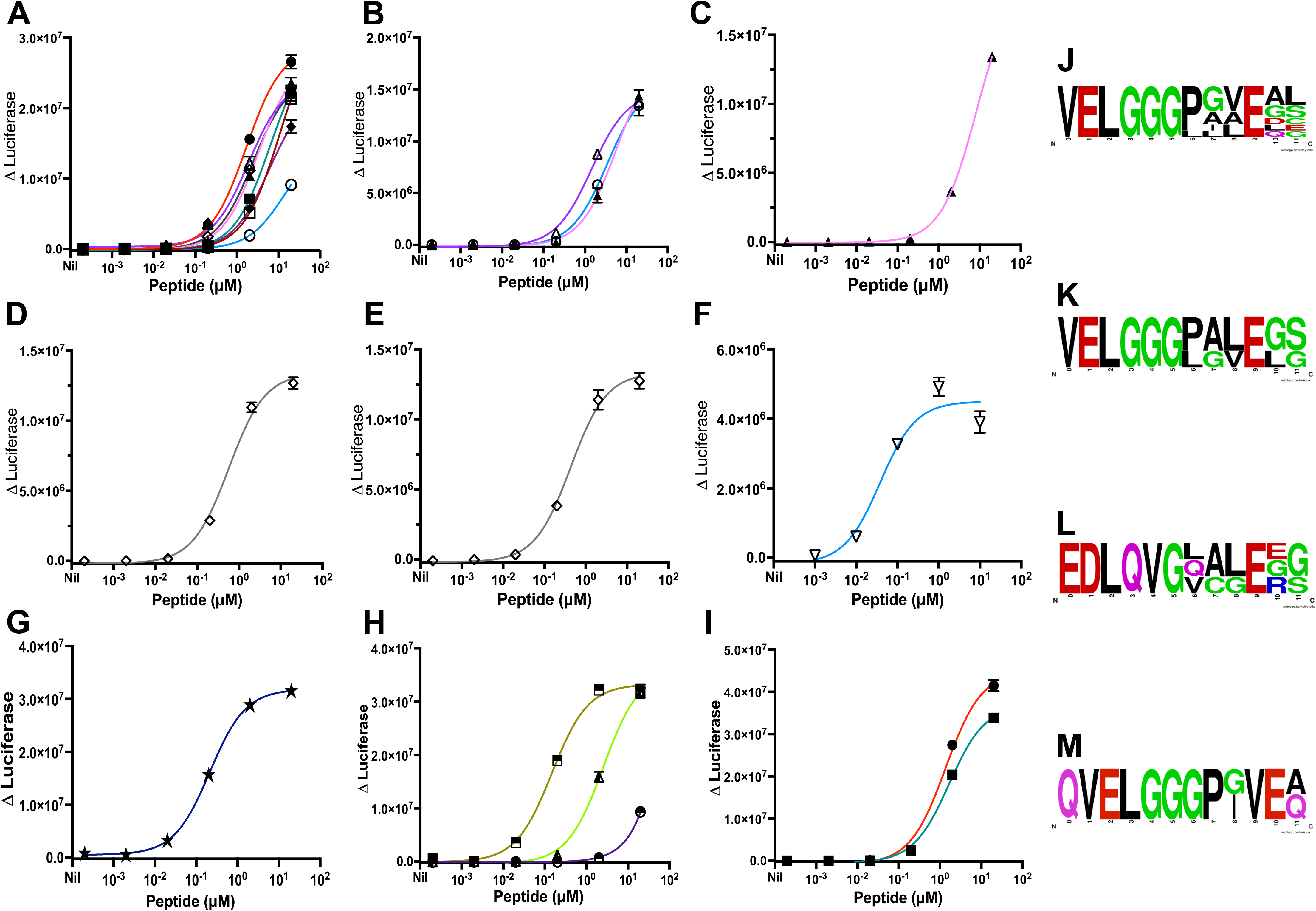
New HIPs are potent T cell agonists. The dose-response curves for the following T-cell avatars are shown: (A) A_4_2 tested against HIP1 (open circles), HIP2 (closed circles), HIP3 (open squares), HIP4 (closed squares), HIP5 (open triangles), HIP6 (closed triangles), HIP7 (open diamonds) and HIP8 (closed diamonds). (B) T cell avatar A_4_5 was tested against HIP1 (open circle), HIP5 (open triangles) and HIP6 (closed triangles). (C) T-cell avatar A_4_6 was tested against HIP6 (closed triangles); (D) T-cell avatar A_4_7 was tested against HIP7 (open diamonds. (E) T-cell avatar A_4_8 was tested against HIP7 (open diamonds). (F) T-cell avatar A_4_28 was tested against HIP9 (inverted triangles). (G) T-cell avatar K_4_143 was tested against HIP7 (open diamonds). (H) T-cell avatar K_4_161 was tested against HIP11 (half shaded circles), HIP12 (half shaded squares) and HIP13 (half shaded triangles). (I) T-cell avatar K_4_207 was tested against HIP2 (closed circles) and HIP4 (closed squares). Antigen-specific responses were measured in triplicate by luciferase assay and represented by Δluciferase. Δluciferase was calculated by subtracting the mean of the ‘no peptide’ treatment from the other treatments. Each point is the mean of triplicate values +/- SD. One representative of at least two experiments is shown. (J-M) show the Logo plots for the epitopes recognized by the T-cell avatars: A_4_2 (J), A_4_5 (K), K_4_161 (L) and K_4_207 (M).

The potency of each HIP varied. The EC_50_ values ranged from 0.02 to >20.00 μM. Importantly, our *E. coli* based screening method revealed HIPs that were weak TCR agonists, for example, HIP 11 which is recognized by the avatar K_4_161 has an EC_50_ of >20.00μM, in contrast, the most potent HIP/avatar, HIP 9/A_4_28 has an EC_50_ of 0.02 μM (Supplementary Table 9). For all T-cell avatars responses were HIP specific. No responses were seen to peptides with native sequences that parallel the HIP sequences (Supplementary Figure 3, Supplementary Table 3). We conclude that the *E. coli* screening method accurately identified new HIPs and most of these HIPs are potent activators of TCRs derived from human islet-infiltrating CD4^+^ T cells.

### Determining the HLA restriction of HIP specific avatars

The HLA restriction of the response to each HIP was determined by blocking with HLA isotype-specific mAbs and testing antigen presenting cells that expressed a single HLA class II allele. As expected, 11 of 13 (85%) HIPs specific T-cell avatars were HLA-DQ8 restricted (Supplementary Figure 4). However, two (A_4_28 and K_4_143) were restricted by HLA-DR4. A summary of the islet-infiltrating T-cell avatars’ responses to the new proinsulin HIPs is shown in Supplementary Table 9. Hence, responses to HIPs were restricted by HLA alleles strongly associated with risk of developing T1D.

### HIP specific T-cell Avatars respond to islet extracts

Because identifying a specific HIP species, or set of HIPs, in human islets is very challenging [28, 29], we took a functional approach to demonstrate the relevance of T-cell specific for the HIPs we have identified. We tested the capacity of HIP specific T-cell avatars to respond to peptide enriched extracts of human islets and spleen which we used a control tissue. Of the nine avatars tested, six (five from Donor A and one from Donor K) responded strongly to islet, but not to spleen extracts (Figure 3). The remaining three avatars had weak, albeit statistically significant, responses to islet peptide extracts compared to the control spleen peptide extracts. This reveals that the T cell avatars respond to peptides present in human islet, but not spleen.

**Figure 3.**
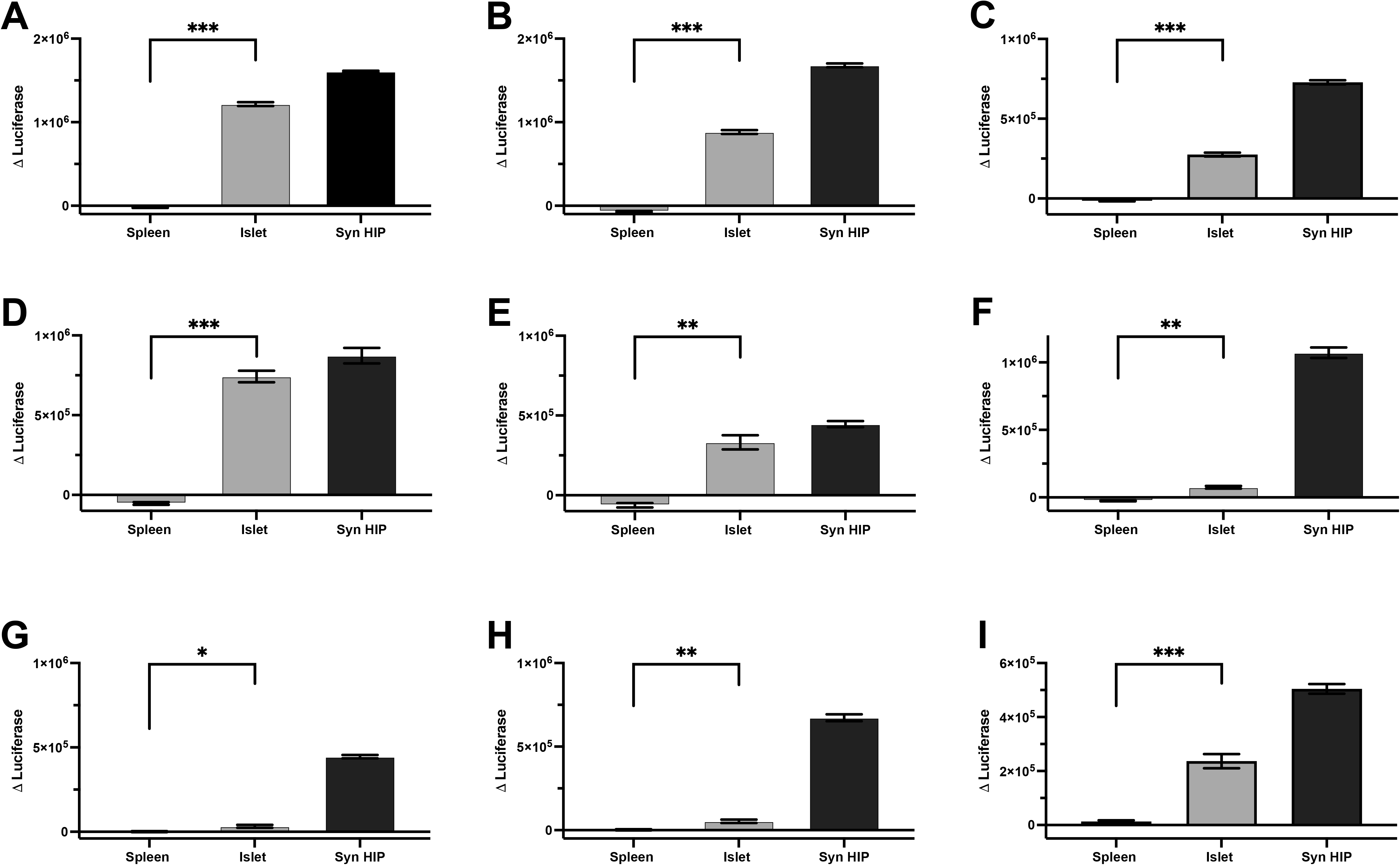
T-cell avatars that recognize new HIPs also respond islet-derived peptides. The following T-cell avatars: A_4_2 (A), A_4_5 (B), A_4_6 (C), A_4_7(D), A_4_8(E), A_4_28 (F), K_4_143 (G), K_4_161 (H), K_4_207 (I) were tested for their capacity to respond to peptide enriched islet and spleen extracts (diluted 1:200 in media). Positive controls were synthetic HIP peptides. Responses were measured by luciferase assay and represented by Δluciferase, calculated by subtracting the mean of 1:200 dilution of DMSO in media treated samples from the other treatment groups. One representative of two experiments is shown. Statistical significance was determined using a paired Student’s t test using two stage linear step up procedure Benjamini, Krieger and Yekutieli statistical hypothesis comparing spleen extract with the islet extract and defined as *p<0.05, **p<0.01, ***p<0.001.

### HIPs stimulate CD4^+^ T-cell responses in the peripheral blood of people with T1D

We used the CFSE-based proliferation assay [30] to determine if responses to the proinsulin HIPs we had identified were detectable in the peripheral blood mononuclear cells (PBMCs) of 10 people with T1D and 10 people without T1D who expressed the HLA-DR4-DQ8 haplotype (Supplementary Tables 7 and 8). Where a T-cell avatar responded to several HIPs, the most potent HIP of this family was used (Supplementary Table 9) for these experiments. A summary of the PBMC responses to seven HIPs is shown in Figure 4. Overall CD4^+^ T-cell responses to HIPs were more frequently detected in PBMC from people with T1D than those without T1D (Figure 4A). The magnitude of the response to HIPs was generally greater in the T1D PBMC compared to the non-T1D PBMCs. However, this difference only reached statistical significance for HIP12. Collectively, responses to the HIP peptides (CDI>3.0) were detected significantly more frequently in PBMC from people with T1D than the HLA-matched non-T1D subjects (Figure 4C). This was most conspicuous for CD4^+^ T-cell responses to HIP 12 which were detected in 60% (6 of 10) of T1D samples, but only 10% (1 of 10) of non-T1D samples (Figure 4B). Responses to HIP 6 were detected in 20% of T1D subjects but were not detected in PBMCs from people without T1D. Hence, we conclude PBMC from people with T1D respond more frequently and strongly to the new HIPs we have identified than the control subjects who do not have T1D.

**Figure 4.**
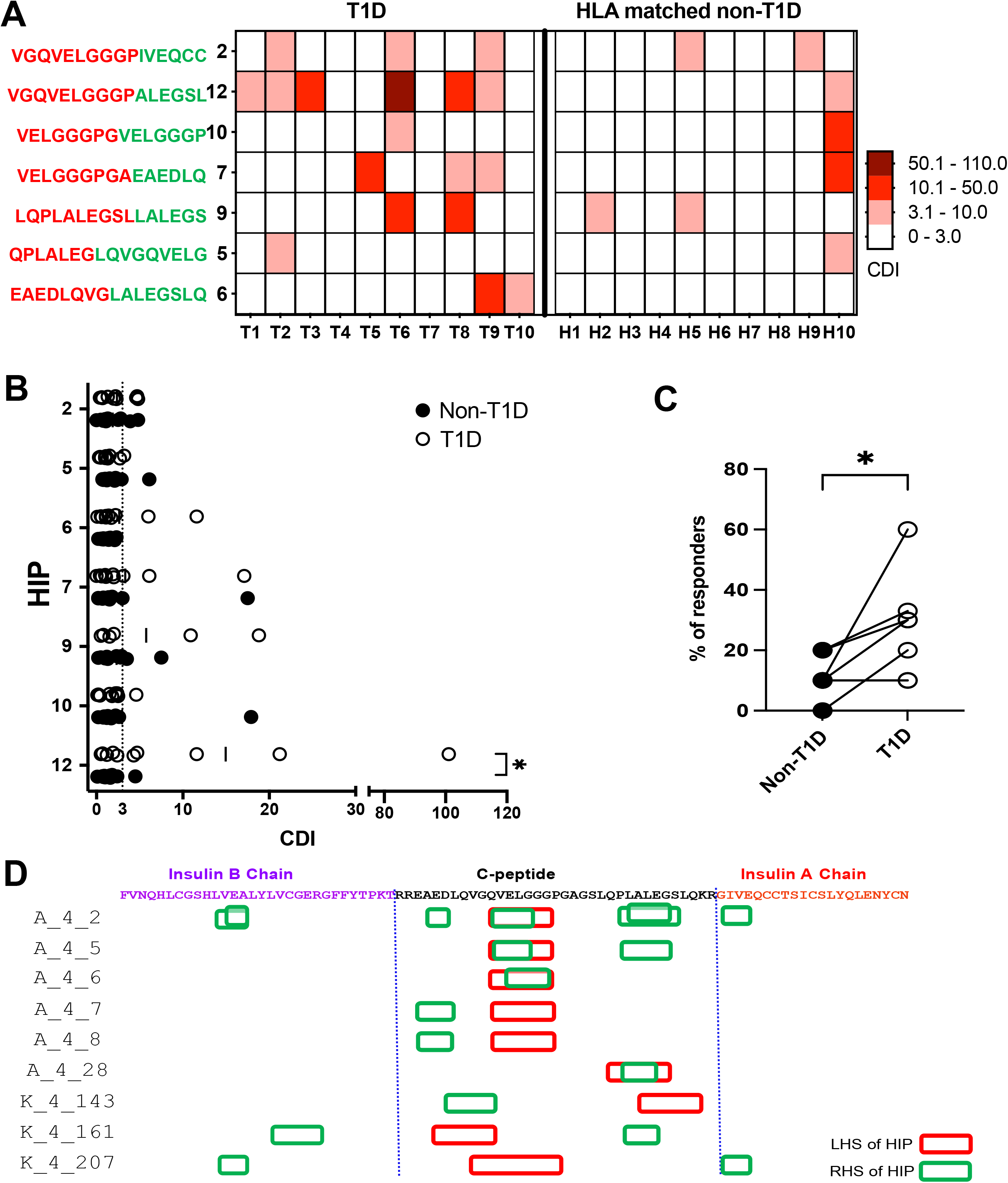
PBMC CD4^+^ T-cells from people with T1D respond to newly identified HIPs. (A) Heat map showing the magnitude of responses to individual HIPs (1 μM), expressed by CDI (cell division index [30]) for PBMC from people with T1D and people without T1D. A CDI <3.0, considered to be a negative response, is represented by a white box. The colour scale indicates the strength of the CD4^+^ T-cell responses to the peptides indicated. ‘X’ indicates not determined (B) The mean CDIs of triplicates for individuals with T1D (n=10) represented by open circle, and without T1D (n=10) HLA matched, represented by closed circle are plotted. Responses with a CDI ≥ 3.0, indicated by the dotted line, are considered to be positive. (C) A comparison of the proportion of the 10 subjects, shown in A and B, that have a detectable CD4^+^ T-cell response to proinsulin HIPs. Statistical significance was determined using unpaired one-tail nonparametric Mann-Whitney test, *p<0.05. (D) Shows a summary of all the HIPs mapped to the location of each fragment in proinsulin. The sequence of proinsulin is shown in a linear form with each amino acid indicated in single letter code. The B-chain is shown in purple, C-peptide in black and A-chain in red text. The C-peptide is bordered by vertical dotted blue lines. The red boxes indicate the NH_2_-terminal peptide fragment and the green the C-terminal peptide fragment. Each line indicates a different T-cell avatar. Avatars from Donor A, start with ‘A_’ and avatars from Donor K start with ‘K_’. Where a T-cell avatar has responded to multiple HIPs there are several boxes shown (for example, avatar A_4_2).

When the origins of each HIP fragment were mapped against the sequence of proinsulin (Figure 4D). it was clear that C-peptide is a major source of peptide fragments which form HIPs. While some sequences appeared in more than one HIP (Supplementary Table 9), all the N-terminal peptide fragments that contribute to HIPs originate from C-peptide. For the C-terminal fragments 8 of the 13 (62%) fragments are derived from C-peptide. In contrast, 3 of 13 (23%) C-terminal fragments are derived from B-chain and 1 of 13 (8%) are derived from A-chain of proinsulin. Hence, we conclude that in addition to being an important antigen in its own right [12], proinsulin C-peptide is a dominant source of peptide fragments which fuse to form HIPs that are recognized by human islet-infiltrating CD4^+^ T cells when presented by HLA-DQ8, or HLA-DR4.

## Discussion

Here we report the identification of 13, HLA-DQ8 or HLA-DR4 restricted, proinsulin HIPs, recognized by TCRs derived from human islet-infiltrating CD4^+^ T cells from two donors who suffered from type 1 diabetes. T cells that recognize these HIPs respond to islet peptide extracts and stimulated CD4^+^ T-cell responses in the PBMC from people with T1D more frequently than those without T1D.

Our optimized screening protocol is ideally suited to the functional identification of new HIPs. The ability to synthesize and clone large libraries allows large numbers of candidate HIPs to be tested. Importantly, we found responses to HIPs across the spectrum of abundance in our library (Supplementary Figure 2B) which suggests that we detected most, if not all, of the possible HIPs. Although the Jurkat avatars are less sensitive to antigen than primary T cells, we still were able to identify HIPs which were relatively weak agonists. In fact, we identified four HIPs that had EC_50_ >10μM (Supplementary Table 9). Which suggests our screen was not unduly biased towards the most potent HIPs. Short peptides are abundantly expressed in *E. coli* and EBV transformed B-cell lines allow us to use autologous antigen presenting cells, relative to the TCRs, to screen the libraries. Our Jurkat reporter line allow rapid screening of many pools and *E. coli* clones and serve as a reliable and robust readout of T-cell recognition. In this study we selected candidate HIPs that were predicted to bind strongly to HLA-DQ8. While 11 of 13 (85%) HIPs specific avatars were DQ8 restricted two were DR4 restricted. Both of these alleles are strongly associated with risk of T1D supporting the relevance of these TCRs and the HIPs they recognize to the immune pathogenesis of T1D. Our approach could easily be modified to select candidate HIPs predicted to bind to any other T1D-associated HLA alleles, such as HLA-DR4, DR3, DQ2, or DQ2/8transdimers.

At the outset, our goal was to make as few assumptions as possible about the HIP sequences. To achieve this, we screened candidate HIPs from the entire sequence of proinsulin and 109 TCRs derived from islet-infiltrating CD4^+^ T cells. Despite this all the HIPs we identified incorporated peptides derived from C-peptide on the N-terminus (Supplementary Table 9). Of the 9 TCR-transduced T-cell avatars that we found to respond in our screen, three were previously shown to respond to C-peptide and two had been shown to recognize a C-peptide-IAPP2 HIP (Supplementary Table 5)[5, 31]. However, previously we had not identified any peptide agonists for the remaining four T-cell avatars, one from Donor A and three from Donor K. Consistent with our earlier work, the N-terminal component of the HIPs recognized by the Donor A T-cell avatars, previously shown to respond to C-peptide derived epitopes, mapped to the central region of C-peptide. Surprisingly, the N-terminal side of the all the HIPs originated from C-peptide, including those recognized by avatars that hadn’t been shown to recognize C-peptide derived epitopes. Most, but not all the C-terminal sequences were also originated from C-peptide (Figure 4D). Two of the HIPs identified here are similar to HIPs reported previously by Delong *et al* [18]. They reported a Cpept-Cpept HIP, (HIP 3: GQVELGGG-EADELQV) which is similar to our HIP7 (VELGGGPGA-EAEDLQ). In our HIP7 there are three amino acids (PGA) more on the N-terminal fragment (Supplementary Table 9). The second pair of similar HIPs are Delong’s HIP 4 (GQVELGGG-GIVEQCC), which is one amino acid different from our HIP 2 (GQVELGGGP-IVEQCC). There is effectively a substitution of P for a G at the junction point.

The reason for strong representation of C-peptide in HIPs is not clear, but several factors may contribute. C-peptide is abundant in the beta cells and unstructured [32-34]. Des-(27-31) C-peptide, a product cathepsin D mediated C-peptide cleavage, is also relatively abundant in islets and may contribute to HIP formation [35-37]. Cathepsin D has broad specificity [38] which is consistent with our findings that there is no clear motif at the points of peptide fusion (Supplementary Table 9). Spontaneous formation of HIPs via an aspartate anhydride intermediate has also been proposed as a mechanism for HIP formation [39]. The HIPs we have identified do not appear to be formed by this mechanism because they do not have the EAED sequence at the N-terminal side of the fusion point.

Similar to others, we found that that some TCRs respond to several HIPs and in some cases cross react with native C-peptide [18, 31, 40]. Some insight into this is gained from the analysis of the TCR/pHLA structures. The structure of TCR/pHLA complexes for three TCRs (A-4_2, A_4_5 and A_4_6) have been solved [31]. This work revealed that the *TRBV5* gene expressed by all these clones facilitated interaction with HLA-DQ8. The N- and C-termini of the HIP interact with TRA and TRB chains respectively. It remains to be determined if the HIPs identified here follow a similar pattern of interaction with HLA-DQ8 and the TCR. Similar cross reactivity has been reported in the NOD mouse, where T cells specific for a HIP comprising insulin B-chain cross react with insulin_B9-23_ [41].

When we examined PBMC responses, we found that each individual with T1D has a CD4^+^ T-cell responses to a unique set of HIPs. This is in concordance with other reports [21, 23, 42], we found that some HIPs more frequently stimulate T-cell responses, but no ‘dominant’ HIP has been identified (Figure 4). This suggests that if HIPs are to be applied to analyse T-cell responses, or as an antigen-specific therapy, in T1D a carefully curated cocktail of HIPs will be required.

There are some limitations of our study. We could only study TCRs derived from four T1D organ donors with widely varying durations of T1D. It is becoming clear that beta-cell antigen specific responses become more difficult to detect as the time since diagnosis with stage 3 T1D passes. Although this is difficult to rigorously demonstrate, without a longitudinal study, it has been our experience. However, we cannot exclude the possibility that HIP specific CD4^+^ T cells are present in the islets of donors with longer-standing T1D that we failed to detect in this study. Our screen was limited to HIPs formed by fusion of peptide fragments derived from proinsulin. Many of our avatars may respond to HIPs formed by the fusion of proinsulin peptide with other beta-cell proteins, but a library of HIPs formed by many beta cell proteins would be unfeasibly large to generate and screen. Our T-cell avatar system allows us to rapidly screen and evaluate TCR specificity, but it is relative insensitive to antigen. Consequently, there may be HIPs in our library that we did not identify because our T-cell readout was not sufficiently sensitive. Nonetheless, we did identify HIPs that were relatively weak agonists (EC_50_> 20μM). While we show that our HIP specific T-cell avatars respond to a peptide extract from human islets, suggesting that the HIPs we have identified are relevant to human T1D, we have not specifically defined the proinsulin HIPs as the agonist species in these extracts. Nonetheless, these lines are known to be very sensitive to the HIPs we’ve identified and the Jurkat avatars did not respond to C-peptide, except in three cases, where we saw very weak responses at the highest concentrations tested (20μM).

In conclusion, we report here 13 new proinsulin derived HIPs. This work greatly expanding our knowledge of repertoire of HIPs recognized by human islet-infiltrating CD4^+^ T cells and places these HIP specific CD4^+^ T cells at the site of autoimmunity in human T1D.

## Materials and Methods

### Subjects and ethical approval

The isolation of pancreatic islets from deceased organ donors was approved by the St. Vincent’s Hospital Human Research Ethics Committee (approval no. SVH HREC-A 011/04). Ethical approval was given by St Vincent’s Hospital HREC (Approval Numbers: 2022/PID05221 and HE135/08) and Southern Health/Royal Children’s Hospital (approval number: 12185B). All participants provided written informed consent. Participants are without T1D or diagnosed according to American Diabetes Association criteria.

### Isolation and TCR sequencing of human islet infiltrating CD4^+^T cells

The isolation of pancreatic islets and the analysis of islet-infiltrating CD4^+^ T cells from Donor A has been reported previously [5]. For the other donors the pancreatic islets were digested by Accutase (Merck) to obtain a single-cell suspension. These cells were stained with a cocktail of mAb conjugated with DNA-barcoded oligonucleotide that are specific for immune cell lineages (TotalSeq™-C Human TBNK Cocktail, BioLegend) according to the manufacturer’s protocol. At the same time, the cells were stained with the following fluorochrome-labelled mAbs: anti-CD45-AF488 (clone QA17A19), and anti-CD3-PE (clone HIT3a). Viable, propidium iodide negative, CD45^+^ and CD3^+^ cells were FACS sorted using FACS Aria Fusion. The scRNA-seq, scTCR-seq and feature barcode libraries of the sorted islet-infiltrating T cells were prepared by using the 10X Genomics Chromium Single Cell Immune Profiling Solution Kit (5⍰ Gene Expression and V(D)J) as per the manufacture’s protocol. The libraries were sequenced following Illumina’s specifications using paired-end sequencing (2 × 150 bp) on Illumina NovaSeq system with a minimum of ∽5,000 reads/cell for V(D)J and feature barcode libraries. The scRNA-seq and scTCR-seq reads were aligned to the 10x pre-built reference genome to GRCh38 (GRCh38-2020-A and vdj_GRCh38_alts_ensembl-5.0.0) and quantified using Cellranger multi pipeline (10x Genomics, version 6.0.1). Filtered gene count matrix from CellRanger was analysed using R (version 4.1.0) and Seurat (version 4.0.5). Cell-specific filtering was performed by retaining cells with RNA features between 200 and 7500, and less than 5% mitochondrial RNA. Centred log ratio transformation across cells by Seurat’s function NormalizeData was used to normalize raw counts of feature barcode. Counts of gene and feature barcodes for CD4 and CD8 were used to identify CD4 single positive cells, further filtering excluded cells with: a single TRAV, a single TRBV, or multiple TRBV genes. This left cells with a single TRBV and one or two TRAV genes. The 50 most abundant TCR clonotypes from these cells were assembled using a custom R script then synthesized and cloned.

### Lentivirus production

For lentivirus-mediated gene transduction, variable regions of TCR genes were expressed and cloned into modified versions of pRRLSIN.cPPT.PGK-GFP.WPRE. The EGFP gene was excised by BamHI/SalI digestion and replaced by human TRAC or TRBC2 with either a PmeI (TRAC) or SbfI (TRBC2) site at the 5’ end of the TCR constant regions. All islet infiltrating TCR variable alpha (TRAV) and TCR beta (TRBV) genes were synthesized by Integrated DNA Technologies (IDT) as gene blocks and cloned into the modified pRRLSIN vectors by In-fusion cloning (Takara) according to the manufacturer’s protocol. Plasmids were extracted from growing *E. coli* and the inserts were verified by Sanger sequencing. For transfection of HEK293T cells, TCR gene-containing plasmids were purified using Macherey-Nagel kits: NucleoBond Xtra Midi Plus (Midi prep), NucleoSpin Plasmid (Standard) or NucleoSpin Plasmid (Transfection Grade). All plasmid purifications were done according to the manufacturer’s recommendations. HEK293T cells were transfected with the appropriate paired TCRs in pRRLSIN and the packaging plasmids: pMDLg/pRRE, pRSV-Rev, and pMD2.g using Lipofectamine 2000 (Invitrogen) as per manufacturer’s instructions. After 7-8 hours incubation at 37°C, 5% CO_2_, the medium was changed. After a further 2.5 days’ incubation, supernatant was collected and filtered through a 0.45μm low protein binding filter. Lentivirus-containing supernatant was either used fresh, or frozen at -80°C until required.

### Generation of T-cell avatars

Starting with Jurkat E6.1, we generated a subline that was TCR deficient, CD4+ and had a luciferase reporter gene knocked immediately following the IL-2 promoter (Supplmentary Figure 1). Briefly, we used CRISPR/Cas9 to generate variant of the human T-cell line Jurkat, that was deficient in TRAC, TRBC and CD4. We then knocked in a NanoLuciferase reporter into the IL-2 locus and isolated a clone that showed an activation induced luciferase response (Supplementary Figure 1). This parental Jurkat lines was then transduced with the TRAV and TRBV constructs that express the TRAV and TRBV genes from human islet-infiltrating CD4^+^ T cells. Transduced cells expressing CD3 and TCR were purified by flow cytometry or magnetic bead enrichment. Briefly, cells were resuspended in viral supernatant at 1–2 x 10^6^/ml with 5μg/ml polybrene. Cells were centrifuged at 1,200 rpm (300 g) for 60min at room temperature, then diluted 1:1 in fresh medium and incubated overnight at 37°C, 5% CO_2_ . Flow cytometry was performed on a Becton Dickinson LSR Fortessa to determine the proportion of CD3 and TCR expressing cells. The following anti-human mAbs were used for FACS staining: CD3-PE (UCHT1, BD Biosciences), and anti-TCRαβ-AF647 (IP26, BioLegend). Cells were stained with the appropriate mAb in PBS 0.1% FBS for 20 minutes on ice, then washed twice. Dead cells were excluded by propidium iodide staining. Compensation settings were determined using single-color controls. TCR and CD3 expressing cells were purified by FACS sorting or using REALease CD3 Microbead Kit (Miltenyi) following the manufacturer’s instructions. All JNL T-cell lines used were >85% CD3^+^/TCR^+^.

### HIP library generation *in silico*

The protein sequence of human proinsulin was fragmented, *in silico*, into 12mer sequences with an overlap of 11 amino acid seq. Each proinsulin 12mers fragment sequence was fused with all other 12mers in both directions to obtain every possible HIP from proinsulin. This proinsulin HIP library was then filtered based on the predicted binding to HLA-DQ8 (HLA-DQA1*03:01; O3:02) using IEDB’s HLA binding prediction algorithm (http://tools.iedb.org/mhcii/). The top 21% of DQ8 binders 12mer HIPs sequence were shortlisted. After filtering out the sequence with duplicate core seq, a total of 4,488 candidate HIPs were left behind (Supplementary figure 2A).

### HIP library cloning and validation

The *E coli* codon-optimized nucleotide sequences of these HIPs were obtained using the GenScript web platform. Each HIP oligo sequence was adjoined with the pKE1 BamH1/Asc1 digested cloning site homology sequence in both termini. The 4,488 candidate HIP oligo sequences containing the cloning homology sequences were divided into 136 pools, where each pool contained 33 unique HIPs (sequences available on request). These single-stranded HIP oligo pools were commercially synthesized by Integrated DNA Technology (IDT). The oligo pools were amplified by PCR and cloned, in frame with GST, into the expression vector pKE-1 [27], by In-fusion cloning. Briefly, the oligonucleotide pools were amplified and duplexed using CloneAmp HiFi PCR Premix (Takara) with HIP Pool forward and reverse primer (Supplementary Table 6). The PCR products were purified using the Macherey-Nagel NucleoSpin column as per manufacturer protocol. The pKE-1 vector was linearized using BamH1/Asc1 (NEB) digestion. All amplified and purified HIP oligonucleotides were cloned into the linearized pKE1 vector by In-fusion cloning (Takara) according to the manufacturer’s protocol. After ligation, the plasmids were transformed into *E coli* Stellar Competent Cells (Takara) and selected with ampicillin (50μg/ml) and kanamycin (10μg/ml) containing Luria Broth. The pool library plasmids were purified using Macherey-Nagel NucleoSpin kits. The pool HIP-expressing bacteria was generated by transforming the purified pool plasmid into the Rosetta strain of *E coli* (Novagen). Four pool HIP-expressing Rosetta were combined to make a ‘super-pool’. In this way, 34 super pool HIP library bacteria were generated from 136 HIP pools. All pools and super-pools of Rosetta were frozen in 70 % glycerol stock for future use.

To validate the HIP library a master pool containing all plasmids in the library was prepared. The master HIP Pool that contains 136 pools was amplified by PCR using Qiagen Taq polymerase with amplicon forward and reverse primer (listed in Supplementary Table 4). The PCR products were gel purified using a Macherey-Nagel NucleoSpin Gel Clean-up kit. Amplicon indexing was done using IDT xGen cfDNA & FFPE DNA library preparation kit and sequencing was performed using a 150-cycle MiSeq platform in the AGRF facility. The abundance of the transcript was determined from the single-end library by using Kallisto (v0.46.1) to generate an index file, followed by using Kallisto quant with an estimated fragment length of 81 bp and an estimated standard deviation of 10 to obtain gene-level normalized transcript per million (TPM).

### Using T-cell avatars to identify novel HIPs

The HIP super-pool bacteria were cultured from frozen glycerol stock in LB media with antibiotics at 37°C with agitation 210 rpm. The overnight HIP-expressing bacteria cultures were diluted to an OD_600_ of 0.06 and grown in 48-well plates in 200μl LB medium with antibiotics until an OD_600_ of 0.5 was reached. Protein expression was induced by adding isopropyl β-D-thiogalactopyranoside (IPTG; 2.0 mM, Sigma). Incubation was continued for another 4h at 37°C in a shaking incubator. All bacteria cultures were then centrifuged at 3,000 rpm for 10 min and the bacterial pellet was resuspended into 300μl of RPMI/5% FCS with gentamycin (Sigma Aldrich) (30 μg/ml) to prevent further bacterial growth. 10 μl of resuspended bacteria from each super pool were cultured with 30,000 autologous EBV-transformed B cells in a 96-well Nunclon Delta white microwell tissue culture plate. After culturing overnight, the T-cell avatars expressing TCRs from human islet-infiltrating CD4^+^ T cells were added (10,000 cells/well). The culture was in a final volume of 150μl of RPMI/5% human serum with gentamycin (30μg/ml). After 22-24h of coculture, the NanoLuciferase activity was measured using the Nano-Glo Luciferase assay (Promega) according to the manufacturer’s instructions. Luminescence was measured on a plate reader (Enspire Perkin Elmer). Responses are reported as the Δluciferase, which is calculated by subtracting the average background luciferase reading measured in the negative control cultures from the responses measured in the other treatments.

HIPs that stimulated the T-cell avatars were identified by subdividing the pools and screening individual colonies. When a colony was identified that stimulated a T-cell avatar the plasmid was extracted and sequenced by Sanger sequencing using the pKE1 sequencing primers (Supplementary Table 4).

### Validation of T-cell responses with synthetic peptide and HLA restriction

Peptides (Genscript) were dissolved in DMSO to 5.0mM, aliquoted and stored at ^-^80°C. A full list of peptides is shown in Supplementary Table 3. The CD4^+^ T-cell avatars’ responses to peptides were measured as described above; except synthetic peptides were used as antigens, instead of *E. coli* and the antigen/EBV cells were not cultured overnight before the T-cell avatars were added. A T cell’s response to antigen was measured as luciferase activity using the Nano-Glo Luciferase assay system (Promega) according to the manufacturer’s instructions (as above). The HLA restriction of the HIP-specific CD4^+^ T-cell clones was determined in a two-step process as described previously [12]. Briefly, first, monoclonal antibodies specific for HLA-DR (clone L243), HLA-DP (clone B7/21) and HLA-DQ (clone SPV-L3) were added to the peptide-stimulated T-cell cultures to a final concentration of 1.0 or 5.0 μg/ml. Second, the HLA alleles were determined using a panel of HLA class II deficient T2 cells that were transfected with HLA-DQA and HLA-DQB alleles indicated in the figure legends. The T-cell avatar’s responses were measured as above. For HLA-DR alleles DR3-DQ2 (IHW09022) or DR4-DQ8 (IHW09031) homozygous lines were compared with autologous EBV.

### Preparation of tissue extracts

Frozen pooled islet (400,000 IEQ) and spleen (4.0g) samples from 8 or 5 healthy organ donors respectively. Tissues were resuspended in 5.0ml of 4M guanidine thiocyanide/1% trifluroacetic acid (TFA) and sonicated on ice for 3.0 minutes. The resultant lysate was passed through 70.0μM cell strainer (Corning) and centrifuged at 1,600rpm for 5min and the supernatant collected. C18 Sep-Pak cartridges (Classic Short cartridge, Waters) were used to enrich the peptides from the tissue lysate. The cartridges were prepared by washing with 3 ml of 99% methanol and 1.0 % TFA, then washed with TFA/salt (0.5 % TFA, 0.5% NaCl) then with 2.0 ml of methanol, TFA, water (30% methanol, 1% TFA in water). The 4.5 ml of cell lysate was loaded into C18 Sep-Pak cartridges. The peptides were eluted in one fraction with 6.0mL methanol/water/TFA (80:19:1, v/v/v). The eluate was lyophilized to dryness and dissolved in 30 μl of DMSO. To test for T-cell avatar responses, T cells and autologous EBV-transformed B cells (both at 10,000 cells/well) were cultured in a 96-well white plate (Nunclon Delta), with either a 1: 200 dilution of enriched peptide, or synthetic HIP peptide (1.0μM). T-cell avatar responses were measured as described above.

### The CFSE-based proliferation assay

The CFSE (5,6-carboxylfluorescein diacetate succinimidyl ester) proliferation assays were performed as described previously [30]. Briefly, PBMC from T1D subjects or non-T1D HLA DQ2 or DQ8 matched donors labelled with 0.1μM CFSE (Life Technologies, Carlsbad, CA) were cultured either with: DMSO (0.02%) or HIPs (1.0 μM), or tetanus toxoid (10LfU/ml). After 7 days of culture the cells were washed in PBS and stained on ice with anti-human CD4-AlexaFluor-647 (clone OKT4, prepared in house). CD4^+^ T-cell proliferation was measured by determining the number of CD4^+^, CFSE^dim^ cells for every 5,000 CD4^+^ CFSE^bright^ cells. The results are presented as a cell division index (CDI) which is the ratio of the number of CD4^+^ cells that have proliferated in the presence of antigen: without antigen [43].

### Statistics and graphs

Graphs were plotted using Prism 10.0.3 and LogoPlots generated (using https://weblogo.berkeley.edu/logo.cgi) Graphing and statistical analysis was done using Prism 10.0.3. Statistical significance was determined using one-way ANOVA and corrected for multiple comparisons using Dunnett’s test. Patient responses were compared using the Mann-Whitney test statistical significance was defined as p<0.05 as shown in the figure legends.

## Acknowledgements

We thank Operational Infrastructure Support Program of the Victorian Government for support. PB is the recipient of a Rising Star Award from St. Vincent’s Institute. This work was funded by the Juvenile Diabetes Research Foundation [2-SRA-2020-909-S-B]. We thank Duncan Campbell for his assistance with the peptide extract preparation. We also thank the Tom Mandel Islet Transplantation Program for islet isolations and the participants and their families for their support to this project.

## References

1. Mannering, S.I., V. Pathiraja, and T.W. Kay, The case for an autoimmune aetiology of type 1 diabetes. Clin Exp Immunol, 2016. 183(1): p. 8–15.

2. Ziegler, A.G., et al., Type 1 Diabetes Prevention: A Goal Dependent on Accepting a Diagnosis of an Asymptomatic Disease. Diabetes, 2016. 65(11): p. 3233–3239.

3. Noble, J.A., Immunogenetics of type 1 diabetes: A comprehensive review. J Autoimmun, 2015. 64: p. 101–12.

4. Noble, J.A. and H.A. Erlich, Genetics of Type 1 Diabetes. Cold Spring Harbor Perspectives in Medicine, 2012. 2(1).

5. Pathiraja, V., et al., Proinsulin-specific, HLA-DQ8, and HLA-DQ8-transdimer-restricted CD4+ T cells infiltrate islets in type 1 diabetes. Diabetes, 2015. 64(1): p. 172–82.

6. Kent, S.C., et al., Deciphering the Pathogenesis of Human Type 1 Diabetes (T1D) by Interrogating T Cells from the “Scene of the Crime”. Curr Diab Rep, 2017. 17(10): p. 95.

7. Babon, J.A., et al., Analysis of self-antigen specificity of islet-infiltrating T cells from human donors with type 1 diabetes. Nat Med, 2016. 22(12): p. 1482–1487.

8. Michels, A.W., et al., Islet-Derived CD4 T Cells Targeting Proinsulin in Human Autoimmune Diabetes. Diabetes, 2017. 66(3): p. 722–734.

9. Anderson, A.M., et al., Human islet T cells are highly reactive to preproinsulin in type 1 diabetes. hProc Natl Acad Sci U S A, 2021. 118(41).

10. Mannering, S.I. and P. Bhattacharjee, Insulin’s other life: an autoantigen in type 1 diabetes. Immunol Cell Biol, 2021. 99(5): p. 448–460.

11. Zhang, L., M. Nakayama, and G.S. Eisenbarth, Insulin as an autoantigen in NOD/human diabetes. Curr Opin Immunol, 2008. 20(1): p. 111–8.

12. So, M., et al., Proinsulin C-peptide is an autoantigen in people with type 1 diabetes. Proc Natl Acad Sci U S A, 2018. 115(42): p. 10732–10737.

13. Rodriguez-Calvo, T., et al., Neoepitopes in Type 1 Diabetes: Etiological Insights, Biomarkers and Therapeutic Targets. Front Immunol, 2021. 12: p. 667989.

14. Mannering, S.I., A.R. Di Carluccio, and C.M. Elso, Neoepitopes: a new take on beta cell autoimmunity in type 1 diabetes. Diabetologia, 2019. 62(3): p. 351–356.

15. Mannering, S.I., et al., The insulin A-chain epitope recognized by human T cells is posttranslationally modified. J Exp Med, 2005. 202(9): p. 1191–7.

16. Mannering, S.I., et al., The A-chain of insulin is a hot-spot for CD4+ T cell epitopes in human type 1 diabetes. Clin Exp Immunol, 2009. 156(2): p. 226–31.

17. Kracht, M.J., et al., Autoimmunity against a defective ribosomal insulin gene product in type 1 diabetes. Nat Med, 2017. 23(4): p. 501–507.

18. Delong, T., et al., Pathogenic CD4 T cells in type 1 diabetes recognize epitopes formed by peptide fusion. Science, 2016. 351(6274): p. 711–714.

19. Haskins, K., Pathogenic T-cell clones in autoimmune diabetes: more lessons from the NOD mouse. Adv Immunol, 2005. 87: p. 123–62.

20. Wiles, T.A., et al., Characterization of Human CD4 T Cells Specific for a C-Peptide/C-Peptide Hybrid Insulin Peptide. Frontiers in Immunology, 2021. 12.

21. Baker, R.L., et al., Hybrid Insulin Peptides Are Autoantigens in Type 1 Diabetes. Diabetes, 2019. 68(9): p. 1830–1840.

22. Baker, R.L., B.L. Jamison, and K. Haskins, Hybrid insulin peptides are neo-epitopes for CD4 T cells in autoimmune diabetes. Curr Opin Endocrinol Diabetes Obes, 2019. 26(4): p. 195–200.

23. Arribas-Layton, D., et al., Hybrid Insulin Peptides Are Recognized by Human T Cells in the Context of DRB1*04:01. Diabetes, 2020. 69(7): p. 1492–1502.

24. Mitchell, A.M., et al., T-cell responses to hybrid insulin peptides prior to type 1 diabetes development. Proc Natl Acad Sci U S A, 2021. 118(6).

25. Wenzlau, J.M., et al., Identification of Autoantibodies to a Hybrid Insulin Peptide in Type 1 Diabetes. Diagnostics (Basel), 2023. 13(17).

26. Mannering, S.I., et al., Identifying New Hybrid Insulin Peptides (HIPs) in Type 1 Diabetes. Front Immunol, 2021. 12: p. 667870.

27. Davis, C.A. and S. Benzer, Generation of cDNA expression libraries enriched for in-frame sequences. Proc Natl Acad Sci U S A, 1997. 94(6): p. 2128–32.

28. Wiles, T.A., L.M. Saba, and T. Delong, Peptide-Spectrum Match Validation with Internal Standards (P-VIS): Internally-Controlled Validation of Mass Spectrometry-Based Peptide Identifications. J Proteome Res, 2021. 20(1): p. 236–249.

29. Wan, X., et al., The MHC-II peptidome of pancreatic islets identifies key features of autoimmune peptides. Nat Immunol, 2020. 21(4): p. 455–463.

30. Mannering, S.I., et al., A sensitive method for detecting proliferation of rare autoantigen-specific human T cells. J Immunol Methods, 2003. 283(1-2): p. 173–83.

31. Tran, M.T., et al., T cell receptor recognition of hybrid insulin peptides bound to HLA-DQ8. Nat Commun, 2021. 12(1): p. 5110.

32. Venugopal, S., et al., Angio-3, a 10-residue peptide derived from human plasminogen kringle 3, suppresses tumor growth in mice via impeding both angiogenesis and vascular permeability. Angiogenesis, 2018. 21(3): p. 653–665.

33. Henriksson, M., et al., Unordered structured of proinsulin C-peptide in aqueous solution and in the presence of lipid vesicles. Cell Mol Life Sci, 2000. 57(2): p. 337–42.

34. Munte, C.E., et al., Solution structure of human proinsulin C-peptide. Febs J, 2005. 272(16): p. 4284–93.

35. Verchere, C.B., et al., Des-(27-31)C-peptide. A novel secretory product of the rat pancreatic beta cell produced by truncation of proinsulin connecting peptide in secretory granules. J Biol Chem, 1996. 271(44): p. 27475–81.

36. Reed, B., et al., Lysosomal cathepsin creates chimeric epitopes for diabetogenic CD4 T cells via transpeptidation. Journal of Experimental Medicine, 2020. 218(2).

37. Crawford, S.A., et al., Cathepsin D Drives the Formation of Hybrid Insulin Peptides Relevant to the Pathogenesis of Type 1 Diabetes. Diabetes, 2022. 71(12): p. 2793–2803.

38. Sun, H., et al., Proteolytic characteristics of cathepsin D related to the recognition and cleavage of its target proteins. PLoS One, 2013. 8(6): p. e65733.

39. Crawford, S.A., et al., Hybrid insulin peptide isomers spontaneously form in pancreatic beta-cells from an aspartic anhydride intermediate. J Biol Chem, 2023. 299(11): p. 105264.

40. Parras, D., et al., Recognition of Multiple Hybrid Insulin Peptides by a Single Highly Diabetogenic T-Cell Receptor. Front Immunol, 2021. 12: p. 737428.

41. Wenzlau, J.M., et al., Insulin B-chain hybrid peptides are agonists for T cells reactive to insulin B:9-23 in autoimmune diabetes. Frontiers in Immunology, 2022. 13.

42. Wiles, T.A. and T. Delong, HIPs and HIP-reactive T cells. Clin Exp Immunol, 2019. 198(3): p. 306–313.

43. Mannering, S.I., et al., An efficient method for cloning human autoantigen-specific T cells. J Immunol Methods, 2005. 298(1-2): p. 83–92.

